# Heterotypic inter-GPCR ß-arrestin coupling regulates lymphatic endothelial junctional architecture in murine lymph nodes

**DOI:** 10.1101/435776

**Authors:** Yu Hisano, Mari Kono, Eric Engelbrecht, Koki Kawakami, Keisuke Yanagida, Andreane Cartier, Sylvain Galvani, Andrew Kuo, Yuki Ono, Satoru Ishida, Junken Aoki, Richard L. Proia, Asuka Inoue, Timothy Hla

## Abstract

Lysophosphatidic acid (LPA) and sphingosine 1-phosphate (S1P) activate G protein-coupled receptors (GPCRs) to regulate key pathobiological processes. Here we report a novel lipid mediator GPCR cross-talk mechanism that modulates lymphatic endothelial junctional architecture in lymph nodes. LPAR1 was identified as an inducer of S1PR1/ ß-arrestin coupling from a genome-wide CRISPR/ Cas9 transcriptional activation screen. LPAR1 activation induced S1PR1 ß-arrestin recruitment while suppressing Gαi protein signaling. Lymphatic endothelial cells from cortical and medullary sinuses of lymph nodes which express LPAR1 and S1PR1, exhibit porous junctional architecture and constitutive S1PR1 coupling to ß-arrestin which was suppressed by the LPAR1 antagonist AM095. In endothelial cells, LPAR1-activation increased trans-endothelial permeability and junctional remodeling from zipper-like structures to puncta of adhesion plaques that terminate at actin-rich stress fibers with abundant intercellular gaps. Cross-talk between LPA and S1P receptors regulates complex junctional architecture of lymphatic sinus endothelial cells, a site of high lymphocyte traffic and lymph flow.

## Introduction

Membrane phospholipids are rapidly metabolized by lipases and synthases to maintain the integrity of biological membranes (1). Lysophospholipids, which are metabolic intermediates, have unique geometry and biophysical properties that facilitate membrane topology, vesicle budding and fusion (2). However, lysophospholipids evolved as extracellular lipid mediators in vertebrates (3). The best characterized are lysophosphatidic acid (LPA) and sphingosine 1-phosphate (S1P), structurally-related lysophospholipids which were originally identified as major regulators of cellular cytoskeletal dynamics (4–6). LPA, which is synthesized in the extracellular environment by autotaxin-mediated hydrolysis of lysophosphatidyl choline, activates six G-protein-coupled receptors (GPCRs) in the EDG and purinergic subfamilies (7). S1P, on the other hand, is synthesized largely in the intracellular environment and secreted via specific transporters SPNS2 and MFSD2B (8–11). Extracellular chaperone-bound S1P activates five GPCRs in the EDG subfamily that are widely expressed (8).

Both LPA and S1P were originally identified as bioactive lipid mediators due to their ability to modulate cytoskeletal dynamics, neurite retraction, cell migration, cell proliferation, and intracellular ion changes (6). Such activity depends on the ability of LPA and S1P to regulate Rho family GTPases (12). After the discovery of the GPCRs for LPA and S1P, genetic loss of function studies in the mice have identified their essential roles in embryonic development and physiological processes of multiple organ systems (13). For example, both LPA and S1P signaling was shown to be important in early vascular development since mice that lack autotaxin (*Enpp2*) as well as sphingosine kinases (*Sphk1* and *2*) were embryonic lethal at early stages of gestation (14–16). Similarly, compound S1P and LPA receptor knockouts also exhibit severe vascular development defects (17, 18). Similar studies have implicated the critical roles of S1P and LPA signaling in neuronal and immune systems (19, 20). A key question that is raised by such findings is whether LPA and S1P are redundant in their biological functions. Data available so far suggest that while some redundant functions are mediated by both LPA and S1P, some unique functions do exist. For example, naïve T cell egress from secondary lymphoid organs is largely dependent on S1P signaling on lymphocyte S1PR1 (21) whereas both LPA and S1P induce fibrotic responses in the lung (22) as well as regulate cardiac development in zebrafish (23). Whether LPA and S1P signaling mechanisms regulate each other (i.e. crosstalk mechanisms) is not known.

The S1PR1 receptor is regulated by molecules that limit its cell surface residency; for example, CD69, GRK2, dynamin, and ApoM+-HDL (24–27). In this report, we searched for novel regulators of S1PR1 coupling to the ß-arrestin pathway. Specifically, we used the TANGO system which uses TEV protease/ ß-arrestin fusion protein and S1PR1-TEV site-tetracycline transcriptional activator (tTA) as a readout (28). Coupled with the single guide (sg)RNA library-directed, CRISPR/ dCas9-induced endogenous genes (29), we screened for novel modulators of S1PR1. The top hit from this unbiased, whole-genome screen was LPAR1. We validated this interaction in a luciferase complementation system that quantifies GPCR coupling to ß-arrestin. Our results suggest that LPAR1 interaction with S1PR1 attenuates S1P signaling in endothelial cells and modulates lymphatic sinus adherens junction and barrier function.

## Results

### Unbiased, genome-wide search for S1PR1 modulators

S1PR1 signaling can be readily monitored by ligand-activated ß-arrestin coupling to the GPCR by the TANGO system, which leads to nuclear fluorescent protein expression (30). This system was shown to be sensitive to receptor activation in transfected cell lines and in the mouse. Since the receptor/ ß-arrestin coupling is faithfully registered and is cumulative due to the stability of the nuclear fluorescent protein, we adapted this system to U2OS osteosarcoma cells that are adaptable to high-throughput screening. Previous work has shown that direct activators of S1PR1, such as CD69 regulate receptor signaling and function (31). In order to search for other endogenous modulators of S1PR1 signaling, we turned to the synergistic activation mediator (SAM) system that uses CRISPR/ Cas9-based, sgRNA-dependent transcriptional activation of endogenous genes (32).

The SAM system turns on endogenous gene expression by sgRNA-dependent recruitment of multiple transcriptional activators (VP64, p65, and HSF1) at upstream of transcription start sites via MS2 bacteriophage coat proteins and mutated Cas9. This screening system was validated by the SAM sgRNA targeting *SPNS2*, an S1P transporter which functions at upstream of S1P receptors (33, 34). The designed SPNS2 SAM sgRNA induced 180-fold increase in its mRNA expression and strongly activated the S1PR1-TANGO signal (Supplemental Figure 1).

To carry out unbiased search for S1PR1-signaling modulators, the SAM sgRNA library was introduced into S1PR1-TANGO system, in which ß-arrestin2 coupling of S1PR1 can be monitored as nuclear expression of Venus fluorescent protein. Venus-positive cells (S1PR1/ ß-arrestin2 signaling positive) were sorted and expanded twice, genomic DNAs were purified and sequenced by Illumina next-gen sequencing (Figure 1A). Bioinformatic analysis indicated that some SAM sgRNA sequences are highly enriched in the Venus-positive cells after sorting (Figure 1B). The *LPAR1* gene was identified as one of the top hits from statistical analysis (Figure 1C). Top ten candidates were individually examined by specific SAM sgRNAs that were enriched after sorting Venus-positive cells. The SAM sgRNA specific for *LPAR1* induced its expression and turned on Venus expression, thus confirming the results from the genome-wide sgRNA screen that identified LPAR1 as an S1PR1 modulator (Supplemental Figure 2).

**Figure 1.**
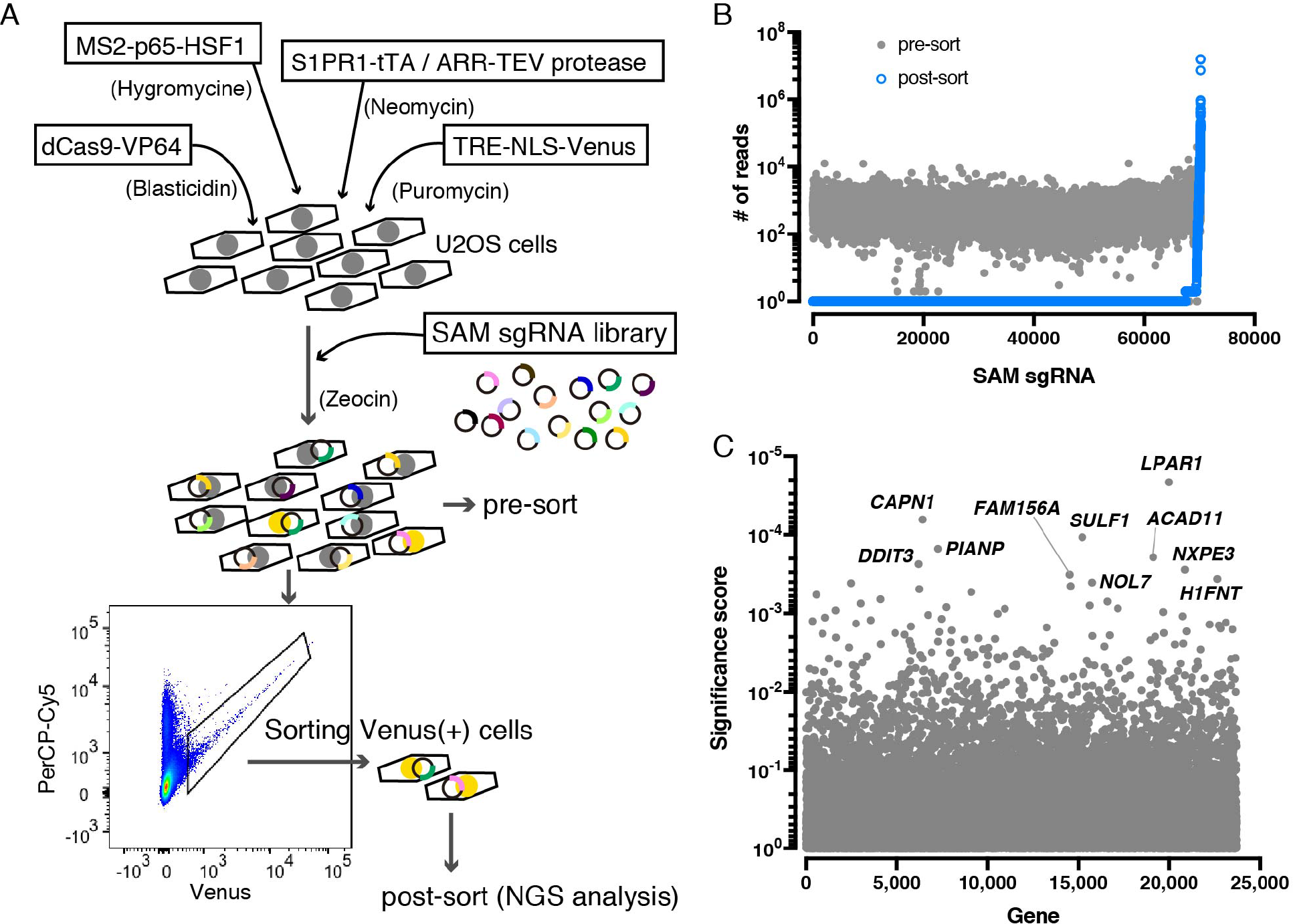
Unbiased whole genome-wide search for S1PR1 modulators Schematic of S1PR1 modulator. (**A**) Schematic of S1PR1 modulator screening system Four lentiviral vectors were transduced into U2OS cell line to enable gene activation by SAM and monitoring S1PR1 activation by TANGO system. The cells introduced with SAM sgRNA library were starved with 0.5% charcoal treated FBS, then the Venus-positive population was sorted and next-gen sequence (NGS) analysis was carried out to identify the enriched SAM sgRNA sequences. (**B**) Scatter plot showing enrichment of sgRNAs after sorting. Most sgRNAs are equally distributed in the pre-sort sample (closed gray circles) while after sorting a small fraction of sgRNAs (2,770 out of 70,290 sgRNAs) were enriched and others were not detected (open blue circles). The y-axis shows the NGS reads of sgRNAs. (**C**) Identification of top candidate genes using the MAGeCK method (68). The names of top ten candidate genes are indicated.

### LPAR1 activation induces β-arrestin recruitment to S1PR1

To further investigate the mechanisms involved in the regulation of S1PR1 signaling by LPAR1, we used the NanoBiT system (35). This system is based on the structural complementation of NanoLuc luciferase and allows one to monitor the protein-protein interactions in real-time. NanoLuc luciferase is split into a small subunit (SmBiT; 11 amino acids) and a large subunit (LgBiT; 18kDa), that are fused with S1PR1 and ß-arrestin1 with mutations in AP-2/ Clathrin-binding motif (to reduce endocytosis), respectively (Figure 2A). S1P dose-dependently stimulated ß-arrestin1 recruitment to S1PR1 in HEK293A cells transfected with S1PR1-SmBiT and LgBiT-ß-arrestin1 (Figure 2B). LPA treatment did not induce ß-arrestin1 recruitment to S1PR1, consistent with the fact that LPA is not a high affinity ligand for S1PR1 (36, 37). However, in cells co-expressing LPAR1 and S1PR1-SmBiT, LPA treatment induced ß-arrestin1 recruitment to S1PR1 with an EC_50_ of ~ 10^−7^ M, which is a physiologically-relevant concentration of LPA (Figure 2C).

**Figure 2.**
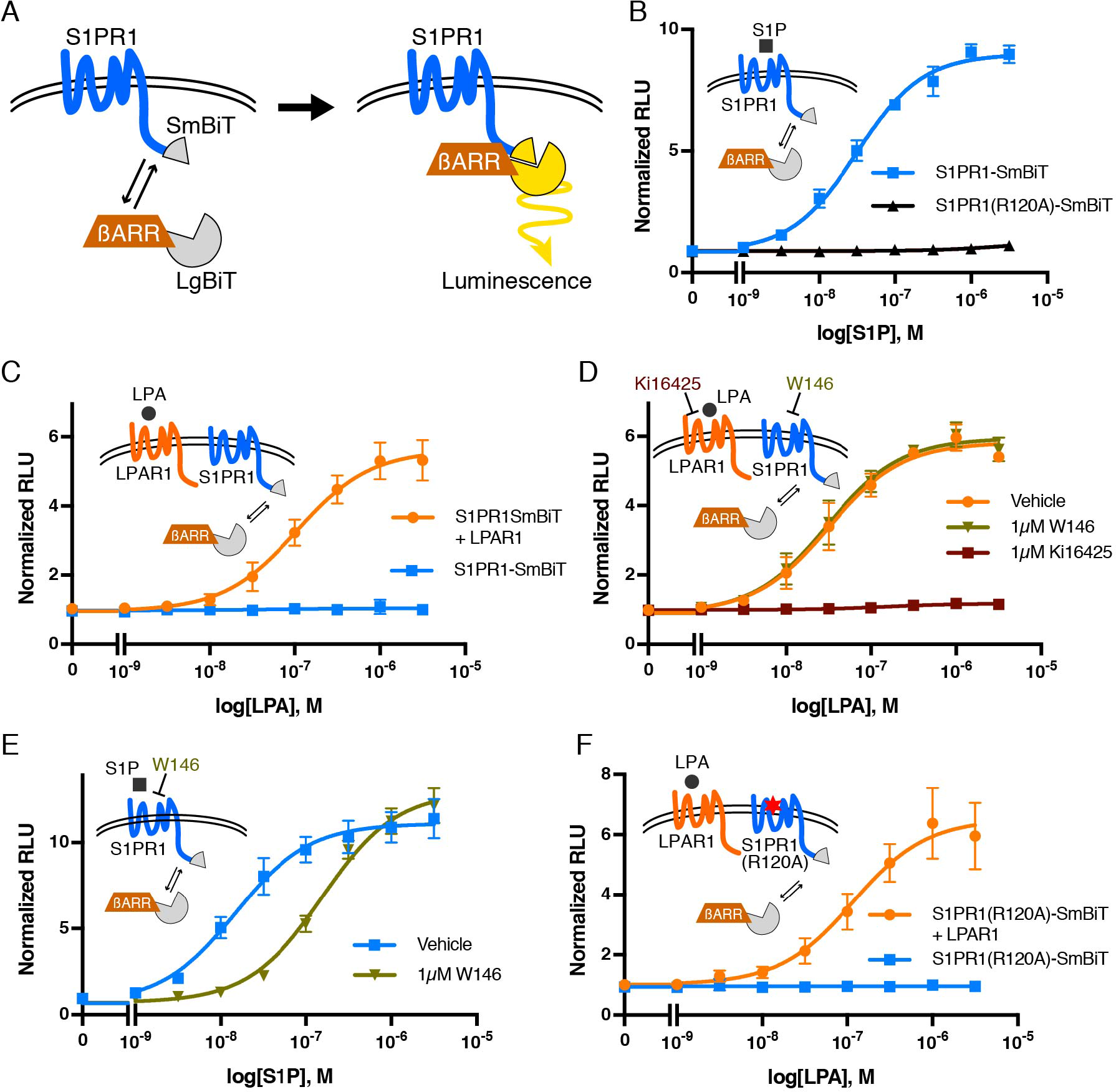
Activated LPAR1 induces S1PR1/ ß-arrestin coupling. (**A**) Schematic of NanoBiT system to measure S1PR1 and ß-arrestin1 interaction SmBiT and LgBiT were fused to C-terminal of S1PR1 and N-terminal of ß-arrestin, respectively. S1PR1 and ß-arrestin1 coupling can be detected as luminescence signal emitted by complementation of SmBiT and LgBiT. (**B**) S1PR1-SmBiT or S1PR1(R120A)-SmBiT was transfected with LgBiT-ß-arrestin1, and luminescence was measured at 15–20 min after S1P stimulation. (**C**) LPAR1 or empty vector was transfected with S1PR1-SmBiT and LgBiT-ß-arrestin1, and luminescence was measured at 15–20 min after LPA stimulation. (**D**, **E**) The cells were incubated with 1 μM Ki16425 or W146 for 30 min prior to stimulation, and luminescence was measured at 15–20 min after LPA (D) or S1P (E) stimulation. (**F**) LPAR1 or empty vector was transfected with S1PR1(R120A)-SmBiT and LgBiT-ß-arrestin1, and luminescence was measured at 15–20 min after LPA stimulation. *n* = 3–8 independent experiments; expressed as mean ± SD.

The effect of LPA was completely blocked by Ki16425, an LPAR1 antagonist(38), indicating that the ß-arrestin1 coupling of S1PR1 is dependent on LPAR1 activation by the ligand (Figure 2D). W146, an S1PR1 antagonist, inhibited S1P-mediated ß-arrestin1 recruitment to S1PR1 but failed to inhibit LPA/ LPAR1-mediated ß-arrestin1 coupling of S1PR1 (Figure 2D and E), suggesting that S1PR1 activation with S1P is not necessary for the LPA/ LPAR1-mediated mechanism to induce S1PR1 coupling to ß-arrestin1. Furthermore, the S1PR1 ligand binding mutant (R120A) behaved similarly to the wild-type S1PR1 by allowing LPAR1 induced ß-arrestin1 coupling (Figure 2B and F). These experiments confirm that LPAR1 activation induced inter-GPCR coupling of ß-arrestin to S1PR1.

### G proteins are not required for LPA/ LPAR1-induced S1PR1/ β-arrestin coupling

LPAR1 couples to three families of G protein alpha subunits (Gαi, Gα12/13, and Gαq/11) while S1PR1 is a Gαi-coupled receptor (39–42). To examine whether LPAR1-induced inter-GPCR coupling of ß-arrestin1 to S1PR1 requires heterotrimeric G proteins, we used HEK293 cells lacking *GNAS*, *GNAL*, *GNAQ*, *GNA11*, *GNA12*, *GNA13*, *GNAI1*, *GNAI2*, *GNAI3*, *GNAO1*, *GNAZ*, *GNAT1*, and *GNAT2* (fullΔGα) generated with CRISPR/ Cas9 system (Supplemental Figure 3 and 4). Even in the HEK293 fullΔGα cells, S1P activation of S1PR1 induced ß-arrestin1 coupling at the same degree with wild-type cells, suggesting that GPCR/ ß-arrestin1 coupling is G protein independent (Figure 2B and 3A), a finding which was reported previously (43). We observed that LPA stimulation of LPAR1 induced S1PR1/ ß-arrestin1 coupling in the HEK293 fullΔGα cells (Figure 3B), indicating that heterotrimeric G protein coupling is not required for inter-GPCR ß-arrestin coupling.

**Figure 3.**
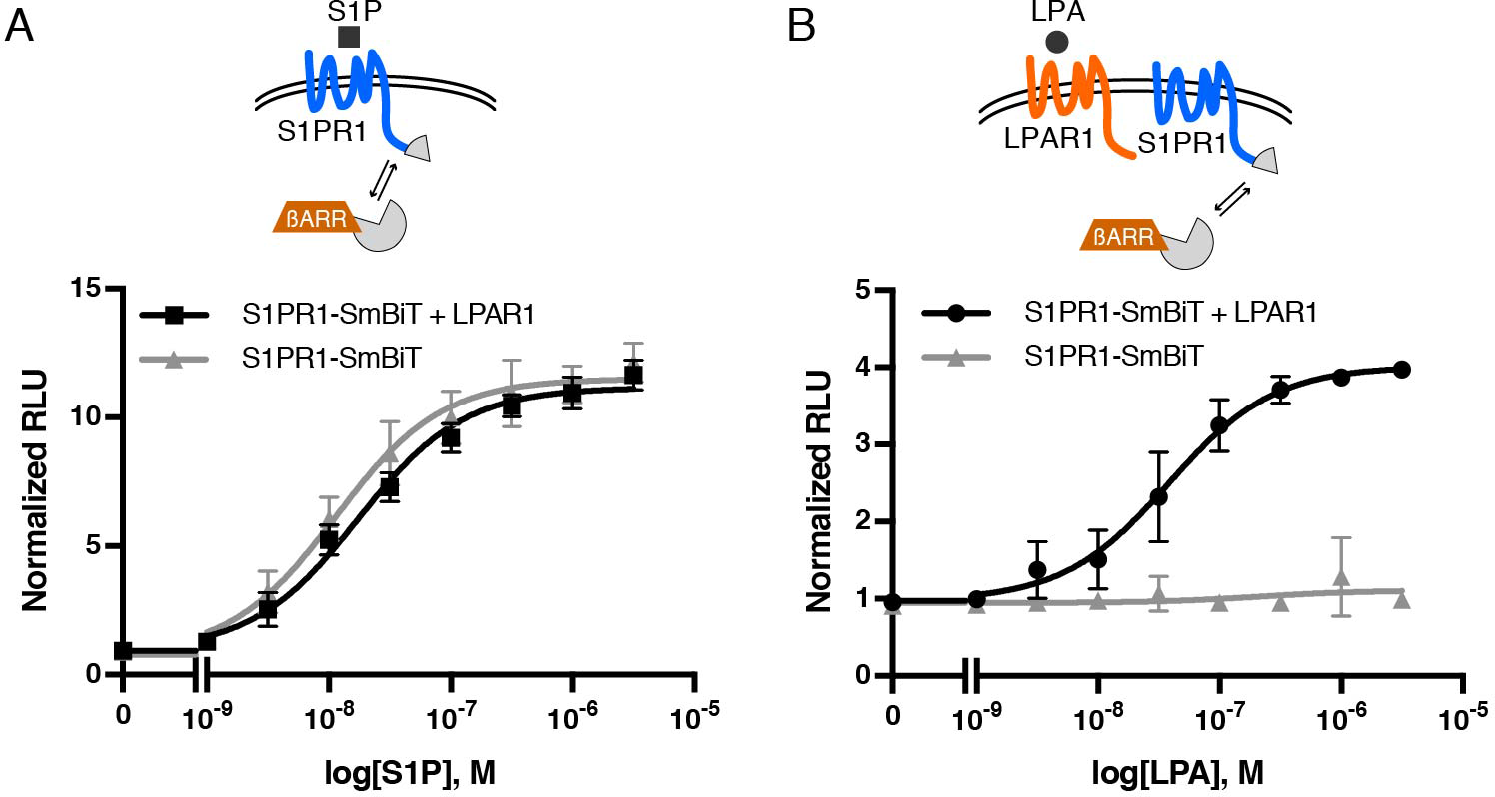
LPAR1-mediated S1PR1/ ß-arrestin coupling in G protein deficient cells. LPAR1 or empty vector was transfected with S1PR1-SmBiT and LgBiT-ß-arrestin1 into HEK293 fullΔGα cells lacking all G protein alpha subunits. Luminescence was measured at 15–20 min after S1P (A) or LPA (B) stimulation. *n* = 3 independent experiments; expressed as mean ± SD.

### LPAR1 C-terminal is necessary for the β-arrestin coupling of S1PR1

ß-arrestin primarily interacts with intracellular C-terminal tail region of GPCRs even though the 3^rd^ intracellular loop may also be involved (44). Deletion of the C-terminal domain in the LPAR1ΔC mutant lost the ability to recruit ß-arrestin1 in response to LPA (Figure 4A) which was demonstrated using the LPAR1ΔC-SmBiT and LgBiT-ß-arrestin1 constructs. Both LPAR1 and LPAR1ΔC mutants couple to the heterotrimeric Gαi protein in an equivalent manner, which was assessed as dissociation of heteromeric G proteins using LgBiT-GNAI2/ SmBiT-GNG (Figure 4B). However, LPAR1ΔC mutant was unable to induce ß-arrestin1 recruitment to S1PR1 in response to LPA (Figure 4C). This result suggests that initial ß-arrestin1 recruitment to LPAR1 is required for the LPA-mediated inter-GPCR coupling of ß-arrestin to S1PR1.

**Figure 4.**
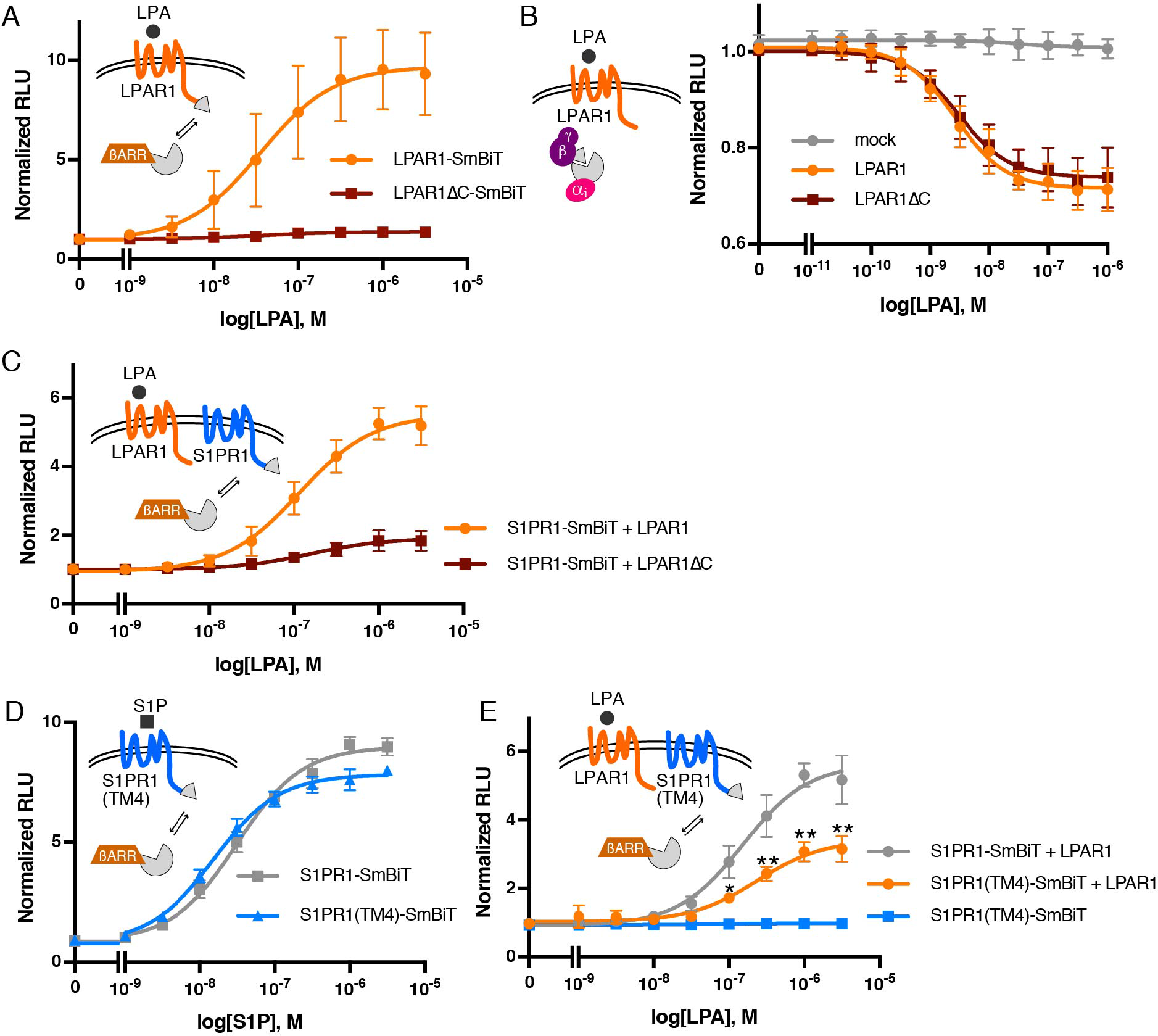
C-terminal of LPAR1 and TM4 of S1PR1 is important for LPAR1-induced inter-GPCR ß-arrestin coupling. (**A**) LPAR1-SmBiT or LPAR1ΔC-SmBiT was transfected with LgBiT-ß-arrestin1, and luminescence was measured at 15–20 min after LPA stimulation. (**B**) G-protein dissociation assay was carried out by transfecting LgBiT-GNAI2, GNB1, and SmBiT-GNGT1 plasmids with LPAR1 or LPAR1ΔC. Luminescence was measured at 6–9 min after LPA stimulation. (**C**)LPAR1 or LPAR1ΔC was transfected with S1PR1-SmBiT and LgBiT-ß-arrestin1, and luminescence was measured at 15–20 min after LPA stimulation. (**D**) S1PR1-SmBiT or S1PR1(TM4)-SmBiT was transfected with LgBiT-ß-arrestin1, and luminescence was measured at 15–20 min after S1P stimulation. (**E**) LPAR1 or empty vector was transfected with S1PR1-SmBiT or S1PR1(TM4)-SmBiT and LgBiT-ß-arrestin1, and luminescence was measured at 15–20 min after LPA stimulation. *n* = 3–5 independent experiments; expressed as mean ± SD. *P* values were determined by two-way ANOVA followed by Sidak’s multiple comparisons test comparing “S1PR1(TM4)-SmBiT + LPAR1” to “S1PR1-SmBiT + LPAR1”; \**P* = 0.0018, \*\**P* ≤ 0.001.

### Transmembrane helix 4 of S1PR1 is important for the β-arrestin coupling of S1PR1

We next examined the hypothesis that direct interactions between S1PR1 and LPAR1 is needed for inter-GPCR ß-arrestin coupling. The transmembrane helix 4 of S1PR1 was reported to interact directly with CD69, a transmembrane C-type lectin (24). The S1PR1(TM4) mutant in which transmembrane helix 4 is replaced with that of S1PR3 decreased the association with CD69, suggesting that it is the domain involved in intermolecular association with GPCR modulators. We therefore, examined the role of the transmembrane helix 4 of S1PR1 in LPAR1-mediated inter-GPCR ß-arrestin coupling to S1PR1. S1PR1(TM4)-SmBiT can be expressed at same level as S1PR1-SmBiT (Supplemental Figure 5) and maintains the ability to recruit ß-arrestin1 by S1P stimulation (Figure 4D). However, the LPAR1-mediated ß-arrestin1 coupling of S1PR1(TM4) was significantly attenuated (Figure 4E), indicating that the transmembrane helix 4 of S1PR1 is important for the LPAR1-mediated ß-arrestin1 coupling of S1PR1.

### LPAR1-induced inter-GPCR ß-arrestin coupling attenuates S1PR1/ Gi signaling

In many GPCRs, ß-arrestin recruitment is an initial trigger for receptor internalization by facilitating interaction with AP-2 and clathrin, that help recruit the GPCRs to the endocytic machinery (45). S1PR1 tagged with Flag at extracellular N-terminal was expressed in HEK293A cells with LPAR1 and Flag-S1PR1 cell surface expression was analyzed by flow cytometry. Surprisingly, Flag-S1PR1 surface expression was not changed by LPA stimulation while S1P stimulation induced Flag-S1PR1 internalization (Figure 5A). Immunofluorescence analysis confirmed these conclusions (Supplemental Figure 6). These results suggest that while LPAR1-induced inter-GPCR ß-arrestin coupling to S1PR1, this event in and of itself is not sufficient to induce S1PR1 endocytosis.

**Figure 5.**
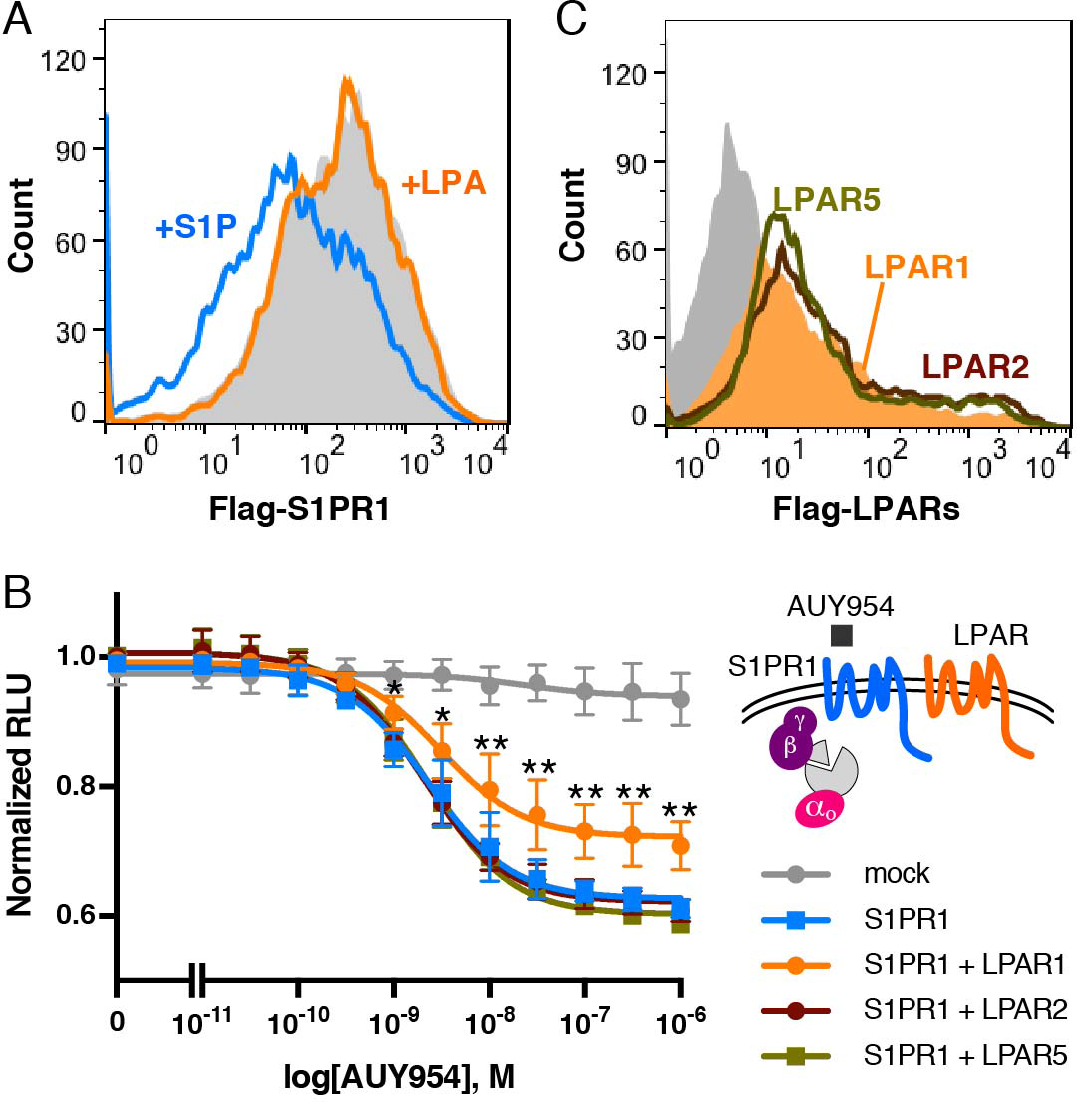
LPAR1 blocks S1PR1/ G protein pathway. (**A**) Flow cytometric analysis showing surface Flag-S1PR1 expression after stimulation with 1 μM S1P (blue line) or LPA (orange line) for 1 hr or without simulation (gray) in HEK293A cells stably expressing Flag-S1PR1 and LPAR1. (**B**) S1PR1 and LPAR1, LPAR2, or LPAR5 were transfected with LgBiT-GNAO1, GNB1, and SmBiT-GNGT1 plasmids. Luminescence was measured at 6–9 min after AUY954 stimulation. *n* = 3–7 independent experiments; expressed as mean ± SD. *P* values were determined by two-way ANOVA followed by Sidak’s multiple comparisons test comparing “S1PR1 + LPAR1” to S1PR1 alone; \**P* ≤ 0.01, \*\**P* ≤ 0.0001. (**C**) Flow cytometric analysis of HEK293A cells transfected with LPAR1 (orange), LPAR2 (brown line), LPAR3 (dark green line) tagged with Flag at N-terminal, or empty vector (gray).

Next, we examined whether LPAR1 activation modulates the S1PR1 signal transduction. Coupling of S1PR1 to the heterotrimeric G protein pathway was assessed using LgBiT-GNAO1/ SmBiT-GNG and AUY954, an S1PR1 selective agonist (46). AUY954 induced S1PR1-mediated heteromeric G protein dissociation in a dose dependent manner, and that was significantly suppressed by co-expression with LPAR1 (Figure 5B). Other LPA receptors (LPAR2 and LPAR5) expressed at similar levels as LPAR1 failed to suppress S1PR1-mediated Gαi protein activation (Figure 5B and C). These results indicate that LPAR1 specifically induces inter-GPCR ß-arrestin coupling to suppress S1PR1 heterotrimeric Gαi protein signaling pathway without inducing receptor endocytosis.

### Endogenous LPAR1 stimulates S1PR1/ β-arrestin coupling in vivo at lymphatic sinuses

Next, to examine whether endogenously-expressed LPAR1 induces inter-GPCR ß-arrestin coupling to S1PR1, we isolated mouse embryonic fibroblast (MEF) cells from S1PR1 luciferase signaling mice, in which endogenous S1PR1/ ß-arrestin2 coupling can be monitored via the firefly split luciferase fragment complementation system (47). As shown in Figure 6A, LPA induced the S1PR1/ ß-arrestin2 coupling in a dose dependent manner, that was blocked by Ki16425, indicating that the activation of endogenously-expressed LPAR1 induces inter-GPCR ß-arrestin coupling to S1PR1.

**Figure 6.**
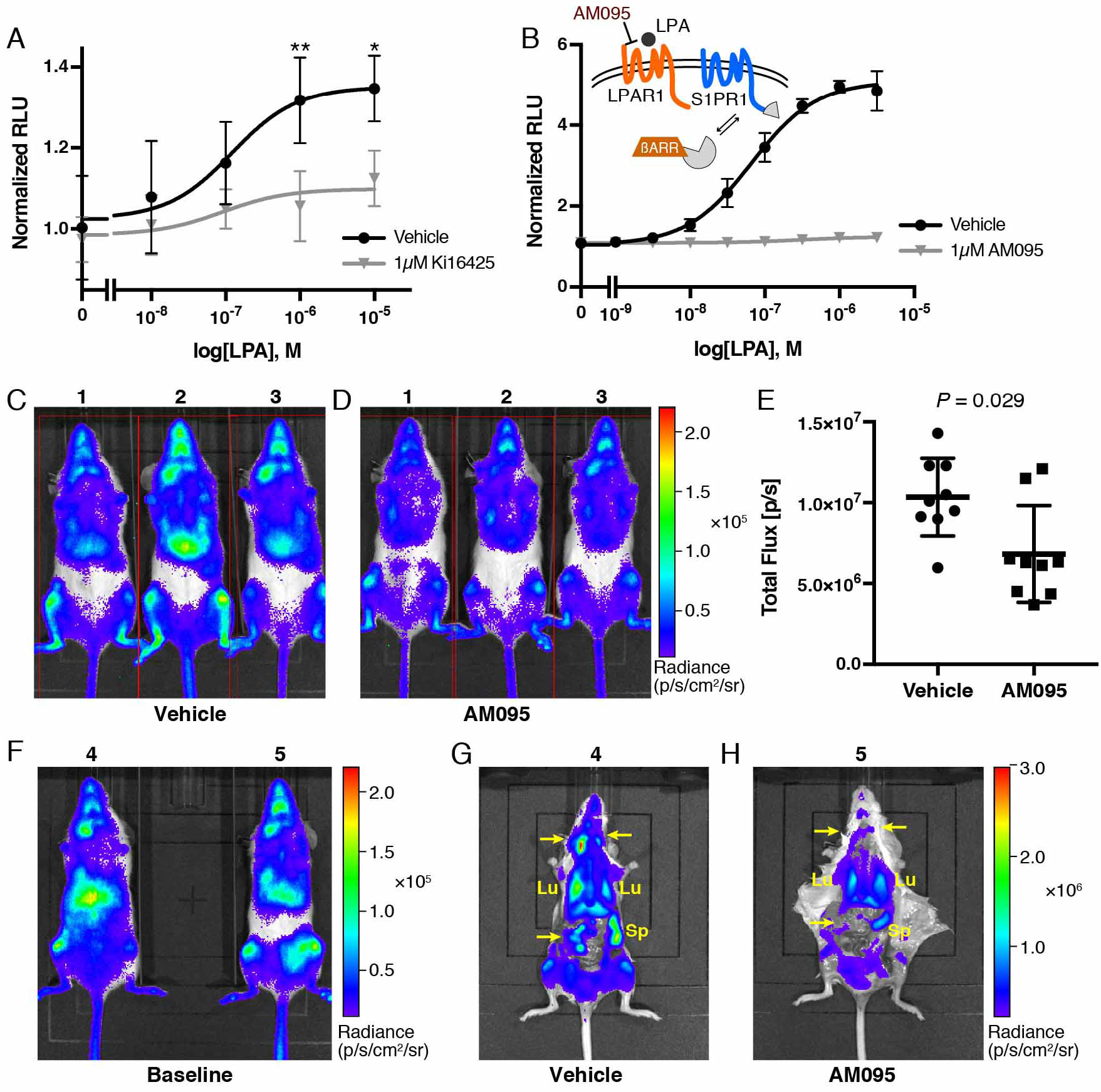
Endogenous LPAR1-induced inter-GPCR ß-arrestin coupling *in vivo*. (**A**) MEF cells isolated from S1PR1 luciferase signaling mice were added with luciferin, then stimulated with LPA at various concentration in the presence or absence of 1 μM Ki16425. Luminescence was measured at 8–12 min after LPA stimulation. *n* = 4 independent experiments; expressed as mean ± SD. *P* values were determined by two-way ANOVA followed by Sidak’s multiple comparisons test comparing vehicle to Ki16425; \**P* = 0.0104, \*\**P* = 0.0021. (**B**) LPAR1 was transfected with S1PR1-SmBiT and LgBiT-ß-arrestin1. The cells were incubated with 1 μM AM095 for 30 min prior to stimulation, and luminescence was measured at 15–20 min after LPA stimulation. (**C**,**D**) Representative bioluminescence images of mice comparing the effects of vehicle (B) or AM095 (30 mg/kg, C), 2 hr after gavage. (**E**) The bioluminescence activity was quantified by determining the total flux (photons/sec; p/s). *n* = 9 for each group; expressed as mean ± SD. *P* value was determined by paired t test. (**F**–**H**) Mice were subjected to imaging prior to administration (E), then dissected in order to image internal organs after vehicle (F) or AM095 (30 mg/kg, G) administration. Arrow, lymph node; Sp, spleen; Lu, lung.

S1PR1 luciferase signaling mice were used to determine if LPAR1-induced inter-GPCR ß-arrestin coupling to S1PR1 occurs *in vivo*. As previously observed, significant S1PR1 coupling to ß-arrestin is seen in several organs in normal mice under homeostatic conditions (47) and (Figure 6C). AM095, an orally available LPAR1 selective antagonist with desirable *in vivo* pharmacokinetic features (48), completely blocked LPA/ LPAR1-mediated ß-arrestin1 coupling of S1PR1 *in vitro* (Figure 6B). Administration of AM095 to S1PR1 luciferase signaling mice significantly decreased bioluminescence signals (Figure 6C–E). Detailed imaging of dissected mice showed that S1PR1 coupling to ß-arrestin in lung, spleen, and lymph nodes were all significantly attenuated by AM095 treatment (Figure 6F–H).

Since lymphatic endothelial cells express both LPAR1 and S1PR1 (49), we further examined the *in vivo* relevance of LPAR1-induced inter-GPCR ß-arrestin coupling to S1PR1 in murine lymph nodes under homeostatic conditions. For this, we used S1PR1-GFP signaling mouse which records cumulative S1PR1 coupling to ß-arrestin while allowing high resolution imaging studies (30). Immunofluorescence and confocal microscopy of brachial lymph node sections in adult mice showed strong S1PR1 coupling to ß-arrestin in lymphatic endothelial cells that make up cortical, medullary, and subcapsular sinuses (Figure 7A). As previously reported (30), high endothelial venules (HEV) also exhibit S1PR1 coupling to ß-arrestin (Supplemental Figure 7). When mice were treated with the LPAR1 inhibitor AM095 for 5 days, S1PR1-GFP signal in subcapsular sinuses and HEV were not altered (Figure 7B,C, and Supplemental Figure 7). In contrast, S1PR1-GFP signal inside the lymph nodes, which are mostly from lymphatic endothelial cells of cortical and medullary sinuses, were suppressed (Figure 7D and E). These data are consistent with quantitative imaging data using S1PR1 luciferase signaling mice shown above and strongly suggest that the site of LPAR1-induced inter-GPCR ß-arrestin coupling to S1PR1 is at the lymphatic endothelial cells of inter-lymphatic sinuses *in vivo*. High resolution images of cell-cell junctions in sinus lining endothelial cells of lymph nodes is shown in Figure 7F. The junctional structure is complex and contains both continuous and punctate VE-cadherin and PECAM-1 positive structures.

**Figure 7.**
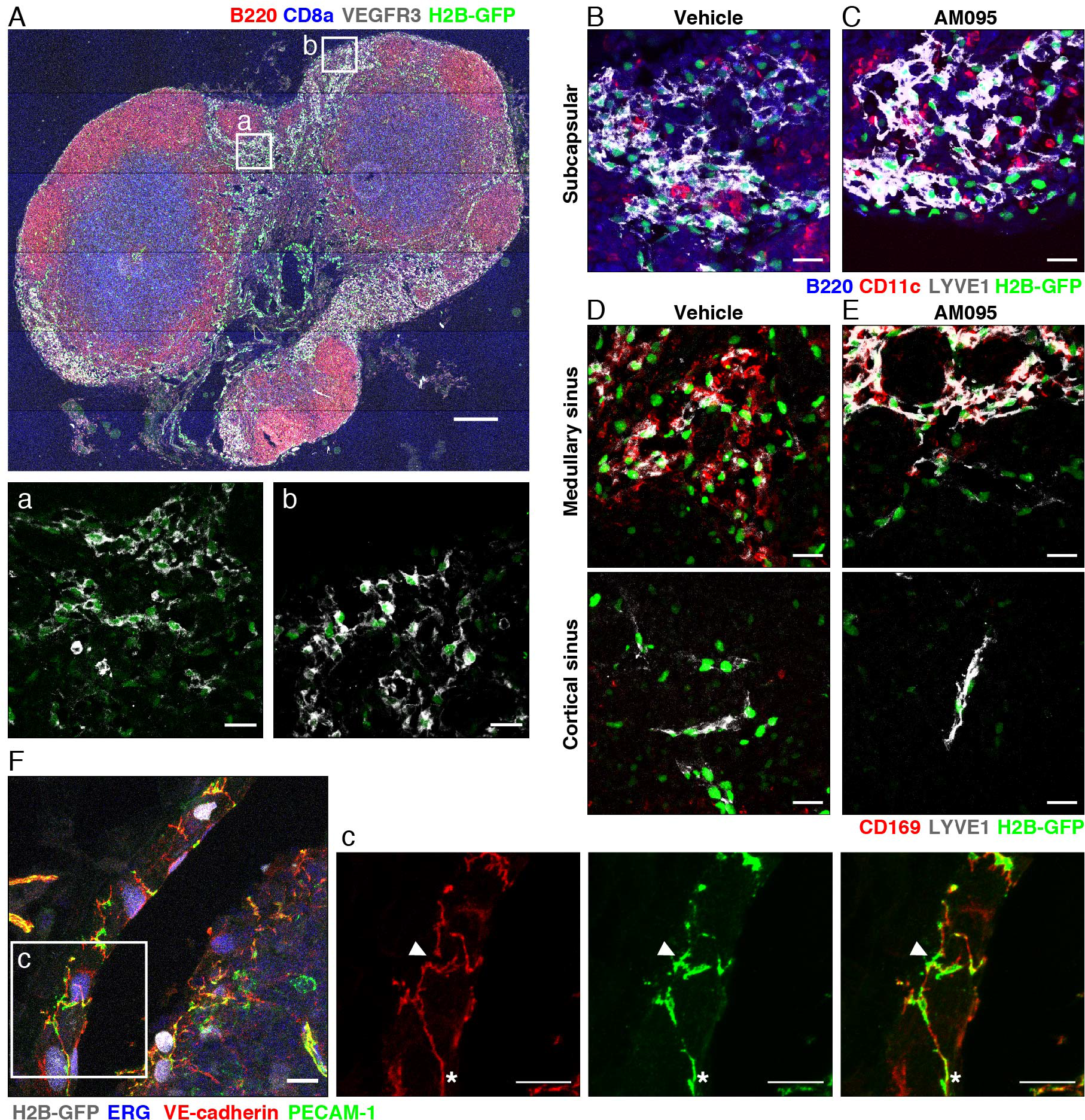
S1PR1/ ß-arrestin coupling in LPAR1 antagonist-treated lymph node. (**A**–**E**) Brachial lymph node sections from S1PR1-GFP signaling mice treated with vehicle or AM095 were stained with B220 (red, B cell), CD8a (blue, T cell), and VEGFR3 (white, LEC) (A), B220 (blue), CD11c (red, dendritic cell), and LYVE1 (white, LEC) (B,C), or CD169 (red, macrophage), and LYVE1 (white) (D,E). LYVE1+ lymphatics were identified as subcapsular sinuses if they were found in subcapsular space and contained B cells and dendritic cells. Medullary sinuses contain CD169+ macrophages, and cortical sinuses are macrophage free (73). (**F**) Mesenteric lymph node section was stained with VE-Cadherin (red), PECAM-1 (green), and ERG (blue). The punctate and continuous junctions were indicated with arrowheads and asterisks, respectively. Bars in (A), (a,b,B–E), and, (F) are 200 μm. 20 μm, and 10 μm, respectively.

### LPAR1 activation suppresses endothelial barrier function and junctional architecture

Lymphatic endothelial cells of sinuses in lymph nodes exhibit complex junctional architecture consisting of button-like structures and high permeability of lymph flow, which are thought to be important for efficient lymphocyte egress and lymphatic fluid drainage and flow (50, 51). Further, endothelial S1PR1 regulates vascular barrier function by activating Gαi/ Rac GTPase signaling pathway that stimulates VE-cadherin assembly at adherens junctions (52). To examine whether LPAR1 modulates S1PR1-dependent barrier function in endothelial cells, LPAR1 was expressed in HUVEC using an inducible system (Supplemental Figure 8) and the barrier function was quantified by measuring trans-endothelial electrical resistance (TEER) (53). As expected, S1PR1 agonist AUY954 induced sustained increase in vascular barrier function (Figure 8A). LPA itself did not influence barrier function either in the presence or absence of AUY954 (Figure 8A). However, in HUVEC expressing LPAR1, LPA-induced a small and transient increase in barrier function (Figure 8C). In sharp contrast, LPA inhibited AUY954-induced vascular barrier increase significantly (Figure 8C). This was completely reversed by Ki16425, an antagonist of LPAR1 (Figure 8D). These data suggest that LPAR1 induces inter-GPCR ß-arrestin coupling to attenuate S1PR1-induced barrier function and thereby enhance the porosity of the endothelial monolayer.

**Figure 8.**
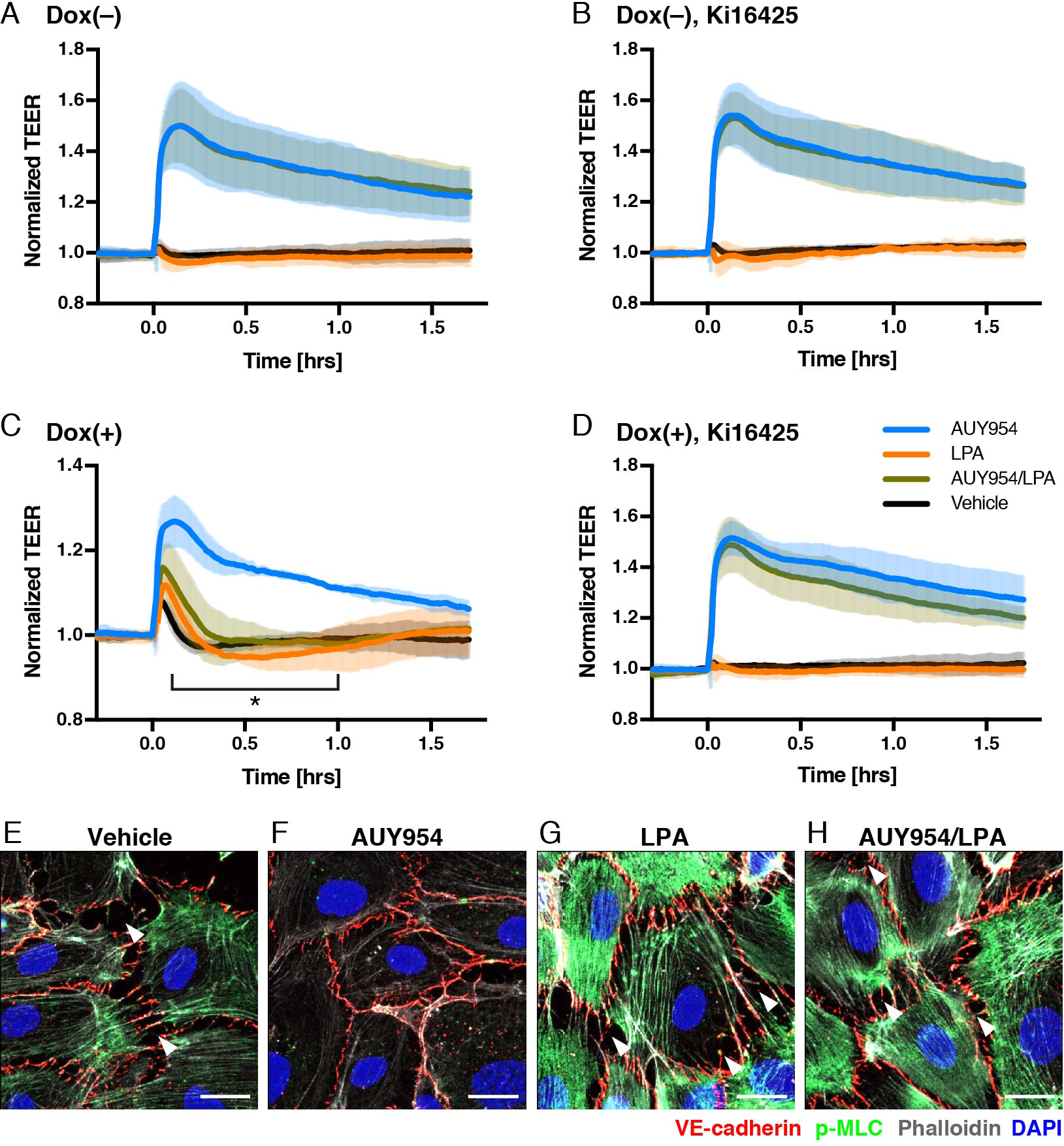
LPA/ LPAR1 attenuates S1PR1-mediated barrier function. (**A**–**D**) HUVECs were analyzed for barrier function by real-time measurement of TEER in the absence (A, B) or presence (C, D) of doxycycline (Dox), which can induce LPAR1 expression by Tet-On system. One day after seeding, the cells were starved with 0.5% charcoal-treated FBS in the absence (A, C) or presence (B, D) of 1 μM Ki16425. At time 0, 100 nM AUY954 (blue), LPA (orange), AUY954 with LPA (dark green), or vehicle (black) was added. *n* = 3 independent experiments; expressed as mean ± SD. *P* values were determined by two-way ANOVA followed by Sidak’s multiple comparisons test comparing “AUY954 + LPA” to AUY954 alone; \**P* ≤ 0.0001. (**E**–**H**) HUVECs expressing LPAR1 were starved with 0.5% charcoal treated FBS for 2 hr, then treated with 100 nM AUY954 and/ or LPA for 30 min. Cells were fixed and stained for VE-Cadherin (red) and p-MLC (green). F-actin and nuclei were stained with phalloidin (white) and DAPI (4′,6-diamidino-2-phenylindole, blue), respectively. Arrowheads indicate intercellular gaps. Bars, 20 μm,

In order to determine the cellular changes induced by LPAR1 and S1PR1 inter-GPCR ß-arrestin coupling, we examined the status of VE-cadherin, a major junctional protein. F-actin and p-MLC (phospho-myosin light chain) were also examined to determine the role of Rho-coupled actin/ myosin architecture which is known to be downstream of LPAR1 (54). As anticipated, S1PR1 activation by AUY954 strongly induced junctional VE-cadherin (Figure 8E and F). In S1PR1 activated HUVEC, minimal intercellular gaps were observed and VE-cadherin appeared as continuous, zipper-like structures at cell-cell borders (Figure 8F). Cortical F-actin was induced and p-MLC staining was attenuated, suggesting increase in Rac GTPase and decrease in Rho GTPase activity, respectively (Figure 8E and F). LPA treatment strongly induced intercellular gaps which punctuate continuous VE-cadherin staining, strong F-actin staining and stress fibers and marked increase in p-MLC staining (Figure 8G). In the presence of both LPA and AUY954, junctional architecture was modulated to contain a hybrid of continuous cell-cell border staining interspersed with punctate VE-cadherin localization at the termini of actin stress fibers (Figure 8H). p-MLC and F-actin at stress fibers was slightly attenuated (Figure 8G and H). However, intercellular gaps were induced when compared with S1PR1 activated HUVEC (Figure 8F and H). These results suggest that the LPAR1 activation induces inter-GPCR ß-arrestin coupling to S1PR1 which modulates Rho GTPase-coupled signal transduction pathways to allow complex cell-cell adherens junction architecture and decreased vascular barrier function. Similar cellular mechanisms may occur in lymphatic endothelial sinuses to regulate high lymphocyte traffic and efficient lymphatic fluid flow.

## DISCUSSION

A major finding of this study is that LPAR1 directly regulates S1PR1 function. This constitutes a heretofore undescribed cross-talk mechanism between LPA and S1P, two lysophospholipids which acquired extracellular functions as vertebrates evolved (3). As vertebrates acquired closed vascular systems, immune cells which are now faced with the challenge of navigating in and out of the circulatory system used S1P, an abundant circulatory lipid mediator with defined spatial gradients for lymphocyte trafficking (21). Our present results suggest that LPA signaling modulates S1PR1 signaling in specific contexts. The S1PR1 receptor is expressed abundantly in endothelial cells and its cell surface expression is controlled by multiple processes (55). For example, the lymphocyte activation-induced molecule CD69 directly interacts with S1PR1 to induce its ligand-dependent endocytosis, a process that decides whether lymphocytes egress occurs or not (24, 31). Indeed, tissue residency of various T cells is controlled by CD69 (31). In endothelial cells, cell surface signaling of S1PR1 regulates vascular barrier function (52, 56). Thus, our finding that LPAR1 modulates S1PR1 directly suggests functional cross-talk between LPA and S1P.

Our study also provides a method to discover novel regulators of GPCR signaling. By adapting a receptor reporter that induces GFP expression downstream of GPCR/ ß-arrestin-coupling with a whole genome-wide CRISPR/ Cas9-dependent transcriptional activation system, we identified LPAR1 as a regulator of S1PR1 function. This system could be adapted to other GPCRs or signaling pathways. Given the modularity and flexibility of CRISPR/ Cas9 system which can both activate or repress genes (29), we suggest that many novel signaling proteins that modulate GPCRs could be identified using similar screens.

We also describe in detail, mechanistic insight into interactions between S1PR1 and LPAR1. S1PR1 and LPAR1 interaction requires the TM4 domain of S1PR1, which was previously identified to be critical for direct interaction with CD69, an event critical for lymphocyte egress (31). Activated LPAR1 recruits ß-arrestin which is then transferred to S1PR1, a phenomenon that we refer to as inter-GPCR ß-arrestin coupling. Recent structural studies indicate that both the C-terminal tail and the 3^rd^ intracellular loop of GPCRs are involved in direct interaction with ß-arrestin (44). Since the 3^rd^ intracellular loop of S1PR1 interacts directly with Gαi family of heterotrimeric G proteins (57), inter-GPCR ß-arrestin signaling resulted in attenuation of S1PR1/ Gαi signaling. However, this mechanism is not sufficient to induce S1PR1 endocytosis. Thus, we suggest that LPAR1-induced inter-GPCR ß-arrestin coupling results in suppression of signaling by plasma membrane-localized S1PR1. This may allow rapid reversal of S1PR1 inhibitory activity and thus may allow acute regulatory mechanism for S1PR1 GPCR.

A key issue we addressed in this study is whether this phenomenon occurs *in vivo*. For this, we turned to the recently-developed real-time S1PR1 luciferase signaling reporter mice, which induces luciferase activity upon S1PR1/ ß-arrestin coupling (47). Our data show that constitutive luciferase signal in the several organs of adult S1PR1 luciferase signaling reporter mice is LPAR1-dependent. In particular, cervical and mesenteric lymph nodes showed strong luciferase activity that was suppressed by LPAR1 antagonist AM095. High resolution confocal microscopy studies show that sinus lining lymphatic endothelial cells in cortical and medullary sinuses of lymph nodes are the cells in which inter-GPCR ß-arrestin coupling between LPAR1 and S1PR1 occurs. Such structures are the sites at which many lymphocytes egress from the lymph node parenchyma into the lumen of the sinuses (50, 51). In addition, lymph from afferent lymphatics that permeate through the lymph node parenchyma flow through these sinus walls to ultimately drain from the efferent lymphatic vessels. We suggest that inter-GPCR ß-arrestin coupling between LPAR1 and S1PR1 regulates the specialized properties of lymph node sinus lining endothelial cells.

It is noteworthy that S1P-dependent lymphocyte egress occurs at cortical and medullary sinuses (58). S1P that is enriched in lymph that is secreted from lymphatic endothelial cells via SPNS2-dependent processes (59, 60), together with low S1P in the lymphatic parenchymal spaces, provides the spatial S1P gradient needed for efficient lymphocyte egress (21). Cell surface S1PR1 on lymphocytes detect this gradient for a spatial cue for the egress process which involves traverse of the lymphocyte through the sinus lining endothelial cells (61). Once the lymphocytes have entered the lumen of the cortical and medullary sinuses, ensuing lymph flow help drain them into efferent lymphatic vessels (58), thus ensuring efficient lymphocyte trafficking. Our results suggest that LPAR1-dependent inter-GPCR ß-arrestin coupling keeps the lymphatic endothelial cell S1PR1 in an inactive state, which may be critical for homeostatic lymphocyte egress. It is noteworthy that LPA is generated in the lymphoid tissue parenchyma (62) and regulate lymphocyte motility and traffic within the lymph node (20, 63).

We addressed the role of LPAR1-induced inter-GPCR ß-arrestin coupling in endothelial cell adherens junctions and barrier function. Our results show that this mechanism alters the junctional architecture and decreases the endothelial barrier function. Specifically, junctions were remodeled from continuous structures at cell-cell borders to punctate structures at the termini of actin-rich stress fibers. This results in the formation of abundant intercellular gaps which explains decreased vascular barrier function. Increased LPAR1-induced Rho GTPase pathways and decreased S1PR1-induced Rac GTPase pathways are likely involved, as determined by the analysis of downstream targets p-MLC and F-actin at the cell cortex and stress fibers, respectively (54, 64).

We propose that junctional remodeling provides a mechanism for high permeability seen in sinus lining lymphatic endothelial cells of lymph nodes. Previous studies in lymphatic endothelial cell junctions have described the presence of button-like junctions which are actively maintained in lymph nodes (51) and in lymphatic capillaries of the small intestinal villi (65). Indeed, this property may allow lymph fluid flow and efficient lymphocyte egress under physiological conditions.

In summary, we have described a mechanism via which LPAR1 suppresses cell surface S1PR1/ Gαi signaling by inter-GPCR ß-arrestin coupling. This process regulates lymphatic endothelial cell junctional architecture and barrier function at sinus lining endothelial cells under physiological conditions. Cross-talk between LPA and S1P receptors regulates complex functions of circulatory and immune systems. Pharmacologic modulation of this pathway may be useful in lymphatic and immune disorders.

## Materials and methods

### Reagents

Primary antibodies used in this study include the following: PE rat monoclonal anti-Flag tag (L5), Alexa Fluor 647 mouse monoclonal anti-HA (16B12), Alexa Fluor 647 rat monoclonal CD8a (53-6.7), Alexa Fluor 647 rat monoclonal CD169 (3D6.112), Alexa Fluor 594 rat monoclonal B220 (RA3-6B2), Alexa Fluore 647 Armenian hamster monoclonal CD11c (N418) (BioLegend); Rabbit polyclonal anti-S1PR1 (H60), mouse monoclonal anti-VE-cadherin (F-8) (Santa Cruz Biotechnology); Rabbit polyclonal anti-phospho-myosin light chain 2 (Cell Signaling Technology); Biotin-conjugated rat monoclonal anti-LYVE1 (ALY7) (eBioscience); Goat polyclonal anti-VEGFR3, Goat polyclonal anti-VE-cadherin (R&D Systems); Rat monoclonal anti-PECAM-1 (MEC13.3) (BD Pharmingen); Rabbit monoclonal anti-ERG (EPR3864) (Abcam). The secondary antibody used for western blotting was HRP-conjugated goat anti-rabbit IgG (Jackson Immuno Research). The secondary antibodies used for immunofluorescence were Alexa Fluor 405 donkey anti-goat IgG (Abcam), Alexa Fluor 647 donkey anti-mouse and anti-goat IgG (Invitrogen), Alexa Fluor 488 goat anti-rabbit IgG (Invitrogen), DyLight 550 donkey anti-rat IgG (Invitrogen), and DyLight 405 donkey anti-rabbit IgG (Jackson Immuno Research). Alexa Fluor 405 streptavidin and Alexa Fluor 546 Phalloidin were from Invitrogen. S1P and LPA were from Avanti Polar Lipids. Ki16425 and AM095 were from Sigma. W146 was from Cayman. AUY954 was from Cellagen Technology.

### Cell culture

HEK293A, HEK293T, and mouse embryonic fibroblast (MEF) cells were cultured in Dulbecco’s modified Eagle’s (DMEM) with L-glutamine, high glucose, and sodium pyruvate medium (Corning) supplemented with 10% fetal bovine serum (FBS) and penicillin-streptomycin (Corning) in a 37 °C incubator with 5% CO_2_. U2OS cells were cultured in McCoy’s 5A medium (Corning) supplemented with 10% FBS and 1% penicillin-streptomycin in a 37 °C incubator with 5% CO_2_. HUVECs were cultured in EGM-2 medium (Lonza) supplemented with 10% FBS or M199 medium (Corning) supplemented with 10% FBS, penicillin-streptomycin, endothelial cell growth factor from sheep brain extract, and 5 units/ml heparin on human fibronectin-coated dishes in a 37 °C incubator with 5% CO_2_.

### Generation of U2OS cell line for library screening

The U2OS cells transduced with dCas9-VP64 (a gift from Feng Zhang, Addgene #61425) and MS2-P65-HSF (a gift from Feng Zhang, Addgene #61426) (32) were selected with 6 μg/mL Blasticidin (Gibco) and 200 μg/mL Hygromycin (Gibco), respectively. For S1PR1-TANGO system, mouse S1pr1 linked to tTA via a TEV protease cleavage site and mouse β-arrestin 2 linked to TEV protease were designed to be cloned in a single vector using a bicistronic internal ribosome entry site (IRES) as described previously (30), and the PCR amplicon from this vector was cloned into pCDH-CMV-MCS-EF1α-Neo lentivector (System Biosciences) with Nhe I and Not I digestion sites. The nuclear localization signal (NLS)-Venus (a gift from Karel Svoboda, Addgene #15753 (66)) with PEST degradation sequence at C-terminal was cloned into downstream of TRE site on pLVX-TetOn lentivector (Clontech). 600 μg/mL Geneticin (G418, Gibco) and 1 μg/mL Puromycin (Gibco) were used for selecting the cells transduced with these constructs.

To produce lentiviral particles, HEK293T cells were seeded on 10 cm dishes 1 day before transfection. On the following day when they had reached 80–90% confluency, medium was replaced by fresh 10% FBS/DMEM medium one hour prior to transfection. 20 μg of lentiviral plasmid, 12.6 μg of pMDL/pRRE, 9.6 μg of pVSV-G, and 6 μg of pRSV-REV were diluted with water and mixed with 85.25 μl of 2M CaCl_2_ solution, then 688 μl of 2 × HBS solution (274 mM NaCl, 1.5 mM Na_2_HPO_4_-7H_2_O, 55 mM HEPES, pH 7.0) was slowly added into the plasmids solution while vortex. After incubation at room temperature for 20 min, the solution mixture was added drop-wise directly to cells. Medium was replaced by 10% FBS/McCoy’s 5A medium 12– 16 hr after transfection. Lentiviral particle-containing supernatant was harvested at 2 days after the medium change, and filtered with a 0.45 μm syringe filter (Corning). PEG-it Virus Precipitation Solution (System Biosciences) was used when concentration was needed. U2OS cells were seeded 1 day before infection. On the following day when they had reached 20–30% confluency, medium was replaced by 10% FBS/McCoy’s 5A medium containing lentiviral particles. Medium was renewed 1 day after infection and antibiotics were added on the following day. The single clones were isolated from antibiotics resistant cells by limiting dilution, then introduced with the SAM sgRNA library (a gift from Feng Zhang, Addgene #1000000057) at a low multiplicity of infection.

### Library screening and sgRNA sequence analysis

The U2OS cells transduced with the SAM sgRNA library were cultured in 400 μg/ml Zeocin (Gibco) to select cells harboring SAM sgRNAs. The Zeocin-resistant cells were allowed to grow (pre-sort cells) or starved with 0.5% charcoal-treated FBS for 2 days. Then, starved cells were harvested and Venus-positive cells were sorted by FACS (post-sort cells) as shown in Figure 1A. The sorted cells were seeded and expanded to repeat sorting. After second expansion, genomic DNAs were harvested from 10 × 107 pre-and post-sort cells using the Quick-gDNA MidiPrep (Zymo Research) according to the manufacturer’s protocol. Amplification and purification of genomic DNAs for NGS analysis was performed as described previously (67). After quality control with Agilent 2200 TapeStation, libraries were subjected to single-end sequencing on an Illumina NextSeq to generate at least 50 million reads for both pre-sort and post-sort cells. Reads were assigned to target genes using the previously described Python script “count_spacers.py” with default parameters (67). The resultant count table was used as input for the script “mageck” to generate significance scores for each target gene (68).

### RNA isolation and quantitative real-time RT-PCR

Total RNA was isolated using TRI reagent (Zymo Research) and further purified by Direct-zol RNA MicroPrep kit (Zymo Research) and treated with DNase (30 U/μg total RNA, QIAGEN) then reverse transcribed using qScript XLT cDNA SuperMix (Quanta Bioscience). Expression of mRNA was quantitated by using PerfeCTa SYBR Green FastMix Reaction Mixes (Quanta Bioscience) and StepOnePlus Real-Time PCR System (Applied Biosystems) with cDNA equivalent to 7.5 ng of total RNA.

Primers used for RT-PCR include the following (5′-3′):

*HPRT*-Fw; TGACACTGGCAAAACAATGCA
*HPRT*-Rv; GGTCCTTTTCACCAGCAAGCT
*SPNS2*-Fw; AACGTGCTCAACTACCTGGAC
*SPNS2*-Rv; ATGAACACTGACTGCAGCAG
*LPAR1*-Fw; ACTGTGGTCATTGTGCTTGG
*LPAR1*-Rv; ACAGCACACGTCTAGAAGTAAC
*FAM156A*-Fw; TATGCTGTTGGGAGGGAAGC
*FAM156A*-Rv; GCAGTATCGACATTCACATCGG

### NanoBiT assay

HEK293A cells were seeded at a density of 8 × 108 cells/6 cm dish 1 day before transfection. On the following day, expression vectors and polyethylenimine (PEI, Polysciences, Inc., pH 7.0) were diluted in 200 μl of Opti-MEM (Gibco), respectively. 300 ng of LgBiT-ß-arrestin1(EE) and 600 ng of GPCR-SmBiT expression vectors were used for ß-arrestin recruitment assay, and 200 ng of LgBiT-GNA, 1000 ng of GNB1, 1000 ng of SmBiT-GNGT1, and 400 ng of GPCR expression vectors were used for G proteins dissociation assay. 10 μl of 1 mg/ml PEI was incubated in Opti-MEM for 5 min at room temperature, then diluted vectors and PEI were combined and mixed with vortex, then incubated for 20 min at room temperature. After incubation, the solution mixture was added drop-wise directly to cells. On the following day, transfected cell were detached with 0.5 mM EDTA/PBS. After centrifugation at 190g for 5 min, cells were suspended in 4 ml of 0.01% fatty acid free BSA (Sigma)/HBSS (Corning) supplemented with 5 mM HEPES (Corning) and seeded on a white 96 well plate at 80 μl/well. 20 μl of 50 μM Coelenterazine (Cayman) was added and incubated for 2 hr at room temperature in dark. Initial luminescence was measured as baseline using SpectraMax L (Molecular Devices), then cells were stimulated with ligands and incubated at room temperature. Luminescence after stimulation was measured and normalized with initial reads. Development and validation of the NanoBiT-G protein dissociation assay is described elsewhere (69).

### Split firefly luciferase complementation assay in MEFs

MEFs isolated from S1PR1 luciferase signaling mice (47) were seed on a white 96 well plate. On the following day, medium was replaced by 80 μl of 0.01% fatty acid free BSA/HBSS supplemented with 5 mM HEPES and incubated for 2 hr at room temperature. 20 μl of 40 mg/mL Luciferin (Perkin Elmer) was added and initial luminescence was measured. After stimulation with LPA, luminescence was measured and normalized with initial reads. Bioluminescence in live mice and internal organs was measured as described previously (47).

### Generation of G protein alpha subunit-depleted HEK293 cells by CRISPR/ Cas9 system

G protein alpha subunit-depleted HEK293 cells were generated by mutating genes encoding members of the Gαi family from previously established HEK293 cells devoid of three Gα families (the Gαs, the Gαq, and the Gα12 families) (43), using CRISPR/ Cas9 system as described previously (70, 71) with minor modifications. sgRNA constructs targeting the *GNAI1*, the *GNAI2*, the *GNAI3*, the *GNAO1*, the *GNAT1*, the *GNAT2*, and the *GNAZ* genes, whose mRNA were expressed in HEK293 cells (72), were designed by a CRISPR design tool (http://crispr.mit.edu) so that a SpCas9-mediated DNA cleavage site (three base pairs upstream of the PAM sequence (NGG)) encompasses a restriction enzyme-recognizing site. Designed sgRNA-targeting sequences including the SpCas9 PAM sequences were as following: 5′-CTTTGGTGACTCAG<u>CCCGGG</u>**CGG**-3′ (*GNAI1*; hereafter, restriction enzyme-site (Sma I in this case) is underlined and the PAM sequence is in bold), 5′-CGTAAAGACCACGGG<u>GATC</u>G**TGG**-3′ (*GNAI2*; Mbo I), 5′-AGCTTGCTTCAGCA<u>GATC</u>CA**GGG**-3′ (*GNAI3*; Mbo I), 5′-AATCGCCTTGCTCCG<u>CTCGA</u>**<u>G</u>GG**-3′ (*GNAO1*; Xho I), 5′-TTTCAGGTGCCGGT<u>GAGTC</u>C**GGG**-3′ (*GNAT1*; Hinf I), 5′-AACCATGCCTCCT<u>GAGCTC</u>G**TGG**-3′ (*GNAT2*; Sac I) and 5′-GATGCGGGTCAGC<u>GAGTC</u>GA**TGG**-3′ (*GNAZ*; Hinf I). The designed sgRNA-targeting sequences were inserted into the Bbs I site of the pSpCas9(BB)-2A-GFP (PX458) vector (a gift from Feng Zhang, Addgene plasmid #42230) using a set of synthesized oligonucleotides as following: 5′-CACC<u>G</u>CTTTGGTGACTCAGCCCGGG-3′ and 5′-AAACCCCGGGCTGAGTCACCAAAG<u>C</u>-3′ (*GNAI1*; note that a guanine nucleotide (G) was introduced at the -21 position of the sgRNA (underlined), which enhances transcription of the sgRNA); 5′-CACC<u>G</u>CGTAAAGACCACGGGGATCG-3′ and 5′-AAACCGATCCCCGTGGTCTTTACG<u>C</u>-3′ (*GNAI2*); 5′-CACC<u>G</u>AGCTTGCTTCAGCAGATCCA-3′ and 5′-AAACTGGATCTGCTGAAGCAAGCT<u>C</u>-3′ (*GNAI3*); 5′-CACC<u>G</u>AATCGCCTTGCTCCGCTCGA-3′ and 5′-AAACTCGAGCGGAGCAAGGCGATT<u>C</u>-3′ (*GNAO1*); 5′-CACC<u>G</u>TTTCAGGTGCCGGTGAGTCC-3′ and 5′-AAACGGACTCACCGGCACCTGAAA<u>C</u>-3′ (*GNAT1*); 5′-CACCGAACCATGCCTCCTGAGCTCG-3′ and 5′-AAACCGAGCTCAGGAGGCATGGTT<u>C</u>-3′ (*GNAT2*); 5′-CACC<u>G</u>ATGCGGGTCAGCGAGTCGA-3′ and 5′-AAACTCGACTCGCTGACCCGCAT<u>C</u>-3′(*GNAZ*). Correctly inserted sgRNA-encoding sequences were verified with a Sanger sequencing (Fasmac, Japan) using a primer 5’-ACTATCATATGCTTACCGTAAC-3’.

To achieve successful selection of all-allele-mutant clone, we performed an iterative CRISPR/ Cas9-mediated mutagenesis. Specifically, in the first round, mutations were introduced in the *GNAZ* gene. In the second round, the *GNAI2*, the *GNAI3*, and the *GNAO1* genes were simultaneously mutated. In the last round, the *GNAI1*, the *GNAT1*, and the *GNAT2* genes were targeted. Briefly, the HEK293 cells devoid of three Gα families (43) were seeded into a 6 well culture plate and incubated for one day before transfection. A plasmid encoding sgRNA and SpCas9-2A-GFP was transfected into the cells using Lipofectamine^®^ 2000 (ThermoFisher) according to a manufacturer’s protocol. Three days later, cells were harvested and processed for isolation of GFP-positive cells (approximately 6% of cells) using a fluorescence-activated cell sorter (SH800, Sony, Japan). After expansion of clonal cell colonies with a limiting dilution method, clones were analyzed for mutations in the targeted genes by a restriction enzyme digestion as described previously (43, 71). PCR primers that amplify the sgRNA-targeting sites were as following: 5’-AGCTGGTTATTCAGAAGAGGAGTG-3’ and 5’-TGGTCCTGATAGTTGACAAGCC-3’ (*GNAI1*); 5′-AAATGGCATGGGAGGGAAGG-3′ and 5′-TAAAACCTCAGTGGGGCTGG-3′ (*GNAI2*); 5′-AGCTGGCAGTGCTGAAGAAG-3′ and 5′-TCATACAAATGACCAAGGGCTC-3′ (*GNAI3*); 5′-GGTCCTTACCGAGCAGGAG-3′ and 5′-CGACATTTTTGTTTCCAGCCC-3′ (*GNAO1*); 5′-TAGGTGTGGCTACGGGGTC-3′ and 5′-GCACTCTTCCAGCGAGTACC-3′ (*GNAT1*); 5’-ACTGCTTCCATCTTAGGTCTTCG-3’ and 5’-CATCAACCCACCCTCTCACC-3’ (*GNAT2*); 5’-CGAAATCAAGCTGCTCCTGC-3’ and 5’-TGTCCTCCAGGTGGTACTCG-3’ (*GNAZ*). Candidate clones that harbored restriction enzyme-resistant PCR fragments were further assessed for their genomic DNA alterations by direct sequencing or TA cloning as described previously (43, 71).

### Measurement of endothelial barrier function *in vitro*

Endothelial barrier function was evaluated by measuring the resistance of a cell-covered electrode by using an endothelial cell impedance system (ECIS) Zθ device (Applied BioPhysics) in accordance with the manufacturer’s instructions. Briefly, arrays were cleaned with 10 mM L-cysteine, washed with sterile water, coated with fibronectin for 30 minutes at 37 °C, and incubated with complete cell culture medium to run electrical stabilization. HUVECs were seeded on a 96 well electrode array (96W10idf) at a density of 2.5 × 104 cells/well in the presence or absence of 1 μg/mL doxycycline. On the following, confluent cells were starved for 2–3 hr in EBM-2 (Lonza) supplemented with 0.5% charcoal treated FBS, then stimulated with AUY954 and/ or LPA. Resistance was monitored and expressed as fractional resistance, normalizing to the baseline at time 0.

### Imaging studies in mice

S1PR1-GFP or luciferase signaling mice have been previously described (30, 47). Bioluminescence image was acquired 2 hr after injection with vehicle (10 μM Na2CO3, 20% 2-Hydroxypropyl-ß-cyclodextrin) through gavage. Three hours after the first imaging for vehicle, the AM095 (30 mg/kg) was administrated to the mice through gavage and bioluminescence image was acquired 2 hr after injection. S1PR1-GFP signaling mice were injected with vehicle or AM095 (20 mg/kg, twice a day) for 5 days through gavage to collect lymph nodes.

### Immunofluorescence staining

HUVECs were washed with cold PBS and fixed with 2% paraformaldehyde (PFA) for 10 min at room temperature. U2OS cell were washed with cold PBS and fixed with cold-methanol for 10 min on ice. Lymph nodes were collected from mice perfused with cold PBS, fixed with 4% PFA, and then embedded in the OCT compound (Sakura Finetek). Cells and cryosections were permeabilized in 0.1% Triton X-100 for 20 min and incubated in blocking solution (75 mM sodium chloride, 18 mM sodium citrate, 1% BSA, 2% FBS, 0.02% sodium azide, and 0.05% Triton X-100) for 1 hr, followed by incubation with primary antibodies for overnight at 4 °C and with secondary antibodies for 2 hr at room temperature. Images were visualized by confocal microscopy using a Zeiss LSM 800. All presented images are 3D reconstructions of z-stack.

### Immunoblot analysis

Cells were washed with cold-PBS and lysed in modified RIPA buffer (50 mM Tris (pH 7.4), 100 mM sodium chloride, 2 mM EDTA, 1% Triton X-100, 0.5% Fos-Choline, and 10 mM sodium azide) containing phosphatase inhibitors (1 mM sodium orthovanadate, 1 mM sodium fluoride, and 5 mM ß-glycerophosphate) and protease inhibitor cocktail (Sigma). After incubation on ice for 30 min and a freeze/thaw cycle, protein concentrations in supernatant from centrifugation at 10,000g, 15 min at 4 °C were determined by bicinchoninic acid assay (Pierce), and denatured for 30 min at room temperature in Laemmli’s sample buffer supplemented with 10% ß-mercaptoethanol. An equal amount of proteins were loaded and separated on an SDS-polyacrylamide gel and transferred electrophoretically to polyvinylidene difluoride membrane (Millipore). Transferred proteins were then probed with rabbit polyclonal anti-S1PR1 (Santa Cruz Biotechnology) and HRP-conjugated goat anti-rabbit IgG (Jackson Immuno Research).

### Flow cytometry analysis

U2OS cells, HEK293A cells, and HUVECs were detached with 0.05% Trypsin (Corning), 0.5 mM EDTA, and Accutase (Innovatice Cell Technologies), respectively. The harvested cells were fixed with 1% PFA for 10 min on ice, and labeled with PE anti-Flag and Alexa Fluor 647 anti-HA antibodies for detecting cell surface expression. The samples were analyzed using BD Calibur FACS system and FlowJo software was used for data analysis.

### Statistical analysis

Data are expressed as means ± SD. Statistical analysis was performed as mentioned using Prism software (GraphPad). *P* values < 0.05 were considered statistically significant.

## Acknowledgments

The authors thank Drs. Hiroko Kishikawa, Yuji Shinjo, and Kumiko Makide for technical assistance of FACS. The SAM sgRNA library, dCas9-VP64, MS2-P65-HSF, and pSpCas9(BB)-2A-GFP (PX458) plasmids were provided by Professor Feng Zhang (Broad Institute of Harvard and MIT). The pCAGGS-ChR2-Venus plasmid was provided by Karel Svoboda.

